# The hippocampus pre-orders movements for skilled action sequences

**DOI:** 10.1101/2024.04.24.590889

**Authors:** Rhys Yewbrey, Katja Kornysheva

## Abstract

Plasticity in the subcortical motor basal ganglia-thalamo-cerebellar network plays a key role in the acquisition and control of long-term memory for new procedural skills, from the formation of population trajectories controlling trained motor skills in the striatum to the adaptation of sensorimotor maps in the cerebellum. However, recent findings demonstrate the involvement of a wider cortical and subcortical brain network in the consolidation and control of well-trained actions, including an area traditionally associated with declarative memory – the hippocampus. Here, we probe which role these subcortical areas play in skilled motor sequence control, from sequence feature selection during planning to their integration during sequence execution. An fMRI dataset collected after participants learnt to produce four finger sequences entirely from memory with high accuracy over several days was examined for both changes in BOLD activity and their informational content in subcortical regions of interest. Although there was a widespread activity increase in effector-related striatal, thalamic and cerebellar regions, the associated activity did not contain information on the motor sequence identity. In contrast, hippocampal activity increased during planning and predicted the order of the upcoming sequence of movements. Our findings show that the hippocampus pre-orders movements for skilled action sequences, thus contributing to the higher-order control of skilled movements. These findings challenge the traditional taxonomy of episodic and procedural memory and carry implications for the rehabilitation of individuals with neurodegenerative disorders.

## Introduction

The neural control of skilled movements is distributed across cortico-striato-cerebellar circuits linked via interlocking loops in the thalamus^1^. As motor skills are learnt, the dominant assumption is that there is a transition from associative-cognitive to motor reference frames^2–4^. This is implemented through a shift towards the involvement of cortical and subcortical areas with mono- and disynaptic projections to brainstem and spinal control centres which control the respective effectors, particularly the contralateral primary motor cortex, the putamen and the ipsilateral anterior lobule of the cerebellum^1,5–7^. This shift is thought to underlie the formation of a motor “repertoire” enabling automatic rather than slow and effortful cognitive control of fine-grained kinematic trajectories and sensorimotor transformations from typing and handwriting to highly skilled musical and athletic performance^8,9^.

Yet, a growing body of evidence from neurophysiology and neuroimaging experiments in both humans and animal models suggests that the network of brain regions engaged in later stages of skill learning and control continues to extend beyond the core motor regions^10^. This applies to skills even after prolonged training over weeks or months^11–15^ and during sequence production entirely from memory without visual or other external guidance^16^. The associated cortical activity patterns have been linked to higher-order control of skilled motor sequences such as the serial order and timing of motor sequences^17^, to enable behavioural flexibility, a key feature of skilled motor control in humans. In contrast, activity within primary motor cortical regions shows a lack of tuning to skilled sequences of movements, even following extensive practice^18–20^.

Whilst the cortical control of motor skills has been characterised in detail over the last three decades, the complementary roles of striato-thalamo-cerebellar loop regions remains poorly delineated. Specifically, it remains unclear whether the encoding of skilled motor sequences, e.g. in striatal and cerebellar effector-related areas is linked to high-order control of sequence chunking and action selection or to lower-order control of movement kinematics^7,21,22^. Furthermore, the hippocampus has long been overlooked as a contributor to procedural motor control due to landmark reports of preserved procedural memory in amnesic patients with medial temporal lobe damage^23,24^. More recently, the original taxonomy has been questioned by findings pointing towards its contribution to sequence learning and consolidation, both during sleep and short breaks^25–29^. Although there is now strong evidence for the hippocampus in procedural motor skill consolidation, its role in the online motor control of skilled movements remains unclear. It remains uncertain whether this contribution is related to the retrieval of task-context associations, or directly tied to the control of specific features of the performed movement sequences.

Here, we set out to tease apart the roles of subcortical areas in skilled movement sequence planning and execution from memory. Participants produced movement sequences with particular order and timing entirely from memory in a delayed sequence production task following two-days of training^16^. Using canonical and multivariate analysis of fMRI patterns in effector-related contralateral striato-thalamo-cerebellar and bilateral hippocampal areas, we probed the tuning of these regions to motor sequences and their ordinal and temporal features during planning and execution. We found that that the hippocampus increased its activity and carried accurate information on the upcoming movement sequence – the movement order independently of the planned sequence timing. In contrast, the activity increase within effector-related striato-thalamo-cerebellar regions during planning and execution was not sequence-specific. This implies their involvement in the lower-order control of individual movement components reused across sequences.

## Results

### Subcortical activity modulation during sequence planning and execution

24 participants were trained to produce four five-finger sequences from memory using their right hand. Behavioural training progressed from production under visual instruction to production from memory across two days. On the final third day, participants produced the sequences entirely from memory in an MRI scanner. The behavioural and cortical results are reported in detail in a previous paper^16^. To isolate neural patterns relating to movement production from those relating to movement preparation, we used two independent trial types: Go trials (Figure 1a), to sample production activity, and No-Go trials (Figure 1b), to sample preparatory activity. Go trials began by cueing the upcoming sequence with an abstract fractal image, followed by a fixation cross, then a black hand with a green background (Go cue) indicated that the sequence should be executed (Figure 1c). A further fixation cross, then performance feedback, followed. No-Go trials were identical to Go trials during movement preparation, however, they did not display a Go cue. Rather, a fixation cross remained on the screen for an extended period. Feedback followed, and participants were rewarded for not producing any presses (see Methods and materials).

**Figure 1.**
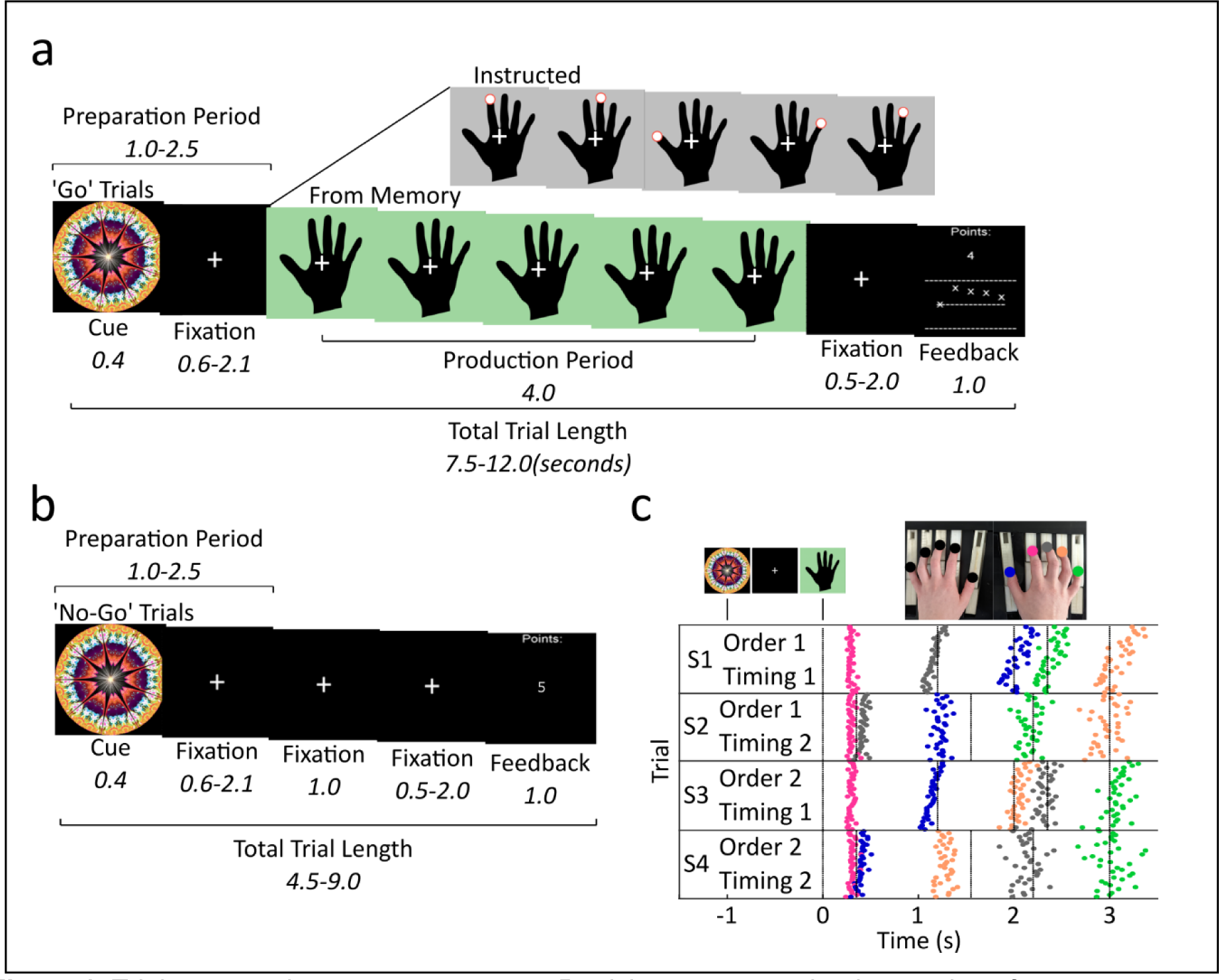
Trial types and target sequences. **a**, Participants were trained to produce four sequences from memory via visual instruction in ‘Go’ trials. The target sequence for each trial was indicated by an abstract fractal cue which preceded a short preparation period. A black hand with a grey (instructed; a red cue indicated which finger to press with a set temporal structure) or green (from memory; participants had to produce the sequence without the visual cue) background appeared to indicate the go cue. A short fixation period followed, after which feedback was provided based on the accuracy and timing of the sequence. **b**, on 50% of trials, the go cue would be replaced by an extended fixation cross. Participants were subsequently rewarded for not making a press during these ‘No-Go’ trials. **c**, all trials for one participant are plotted with the colour indicating which button was pressed according to the image above the plot. Target sequences consisted of permutations of two timings and two orders, constituting S1 through S4. Participants performed the sequences using the right hand on a 10-finger force transducer.

Firstly, we examined whether the subcortical regions of interest with connections to the active effector (right hand) – the left thalamus, caudate, putamen, and the right CB lobules IV and V) – as well as the bilateral hippocampi, changed overall BOLD activity during the delayed sequence production task. We defined all regions of interest (ROI) using each participant’s individual anatomy and extracted the percent signal change during preparation (No-Go trials) and production (Go trials) relative to rest (see Methods and materials). The extracted percent signal change values from across all voxels belonging to each ROI were then averaged for each participant (Figure 2a, b). All tests were Bonferroni corrected twice to account for phase (preparation and production) in each pre-defined ROI. During preparation, we observed activity above baseline in the caudate (*t*(23) = 3.28, *p* = .006, *d* = 0.67), putamen (*t*(23) = 2.67, *p* = .028, *d* = 0.55), and hippocampus on both the left (*t*(23) = 6.28, *p* < .001, *d* = 1.28) and right (*t*(23) = 4.63, *p* < .001, *d* = 0.94) sides. No significant difference relative to baseline was found in the thalamus nor in cerebellar lobules IV and V during preparation (*p* > .114, *d* < 0.41). During production, activity increases were observed in all subcortical area with connections to the active effector - contralateral thalamus (*t*(23) = 7.54, *p* < .001, *d* = 1.54), caudate (*t*(23) = 2.66, *p* = .028, *d* = 0.54), putamen (*t*(23) = 4.21, *p* < .001, *d* = 0.86), cerebellar lobule IV (*t*(23) = 5.61, *p* < .001, *d* = 1.15), and CB lobule V (*t*(23) = 12.22, *p* < .001, *d* = 2.49). The hippocampus, however, showed activity decreases during production on the left (*t*(23) = 2.45, *p* = .044, *d* = 0.50) and right (*t*(23) = 3.19, *p* = .008, *d* = 0.65) sides. In sum, only the striatum and hippocampus were active during sequence preparation from memory. In contrast, the striato-thalamo-cerebellar regions with direct relevance to the active effector increased their activity during production, whereas hippocampal activity decreased.

**Figure 2.**
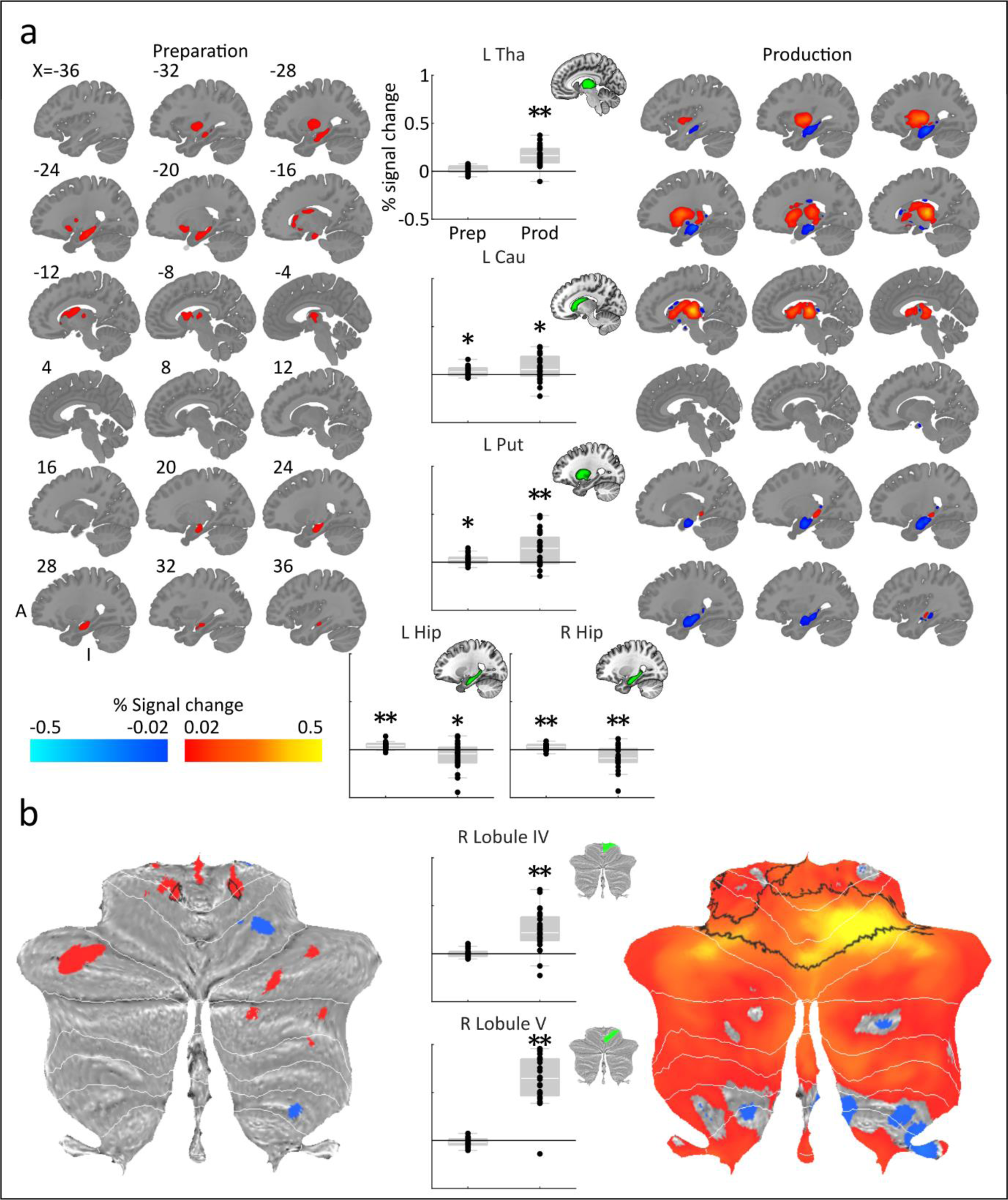
Percent signal change in subcortical regions during preparation and production. **a**, volumetric slices of each subcortical region displaying percent signal change for preparation and production, far left and right respectively. Centre shows the level of percent signal change when averaged across each subcortical region. ** *p* < 0.01; * *p* < 0.05; two-sided t test against 0. Error bars represent standard error of the sample mean. **b**, as above, for cerebellum. Black outline indicates significant clusters. Tha, thalamus; Cau, caudate nucleus; Put, putamen; Hip, hippocampus.

We then localised the peak activations within each subcortical area and across the whole cerebellar volume by identifying significant clusters of activation using a random effects analysis (Figure 2a, b). All results were corrected using SPM’s small-volume correction and Bonferroni corrected twice to account for phase. During preparation, we found significant clusters of activation in the caudate’s ventral anterior head (*t*(23) = 6.02, *pFWE* = .008, extent = 101, MNI coordinates: x=-16, y=22, z=-5), anterior body (*t*(23) = 5.56, *pFWE* = .020, extent = 83, x −16 y 22 z 7), and dorsal anterior tail (*t*(23) = 5.55, *pFWE* = .008, extent =106, MNI coordinates: x=-12, y=-6, z=17), posterior putamen (*t*(23) = 5.71, *pFWE* < .001, extent = 228, MNI coordinates: x=-32, y=-14, z=9), anterior left hippocampus (*t*(23) = 7.97, *pFWE* < .001, extent = 766, MNI coordinates: x=-26, y=-22, z=-15), anterior right hippocampus(*t*(23) = 6.33, *pFWE* < .001, extent = 471, MNI coordinates: x=38, y=-20, z=-17), and CB right lobule V extending into lobule IV (*t*(23) = 5.09, *pFWE* = .002, extent = 205, MNI coordinates: x=17, y=-45, z=17). During production, we found significant clusters of activation widespread in the thalamus which centred middle posterior (*t*(23) = 14.29, *pFWE* < .001, extent = 851, MNI coordinates: x=-16, y=-24, z=5), the anterior tail of the caudate (*t*(23) = 5.24, *pFWE* < .001, extent = 208, MNI coordinates: x=-10, y=0, z=9), posterior putamen (*t*(23) = 7.06, *pFWE* < .001, extent = 783, MNI coordinates: x=-28, y=-18, z=3), and CB lobules IV-VI (the peak of which is in lobule VI, outside of our pre-defined cerebellar ROIs; *t*(23) = 8.46, *pFWE* < .001, extent = 20079, MNI coordinates: x=7, y=-69, z=-16). The location of the clusters generally shifted or expanded from associative to motor subregions of the subcortical areas of interest upon execution.

### Hippocampal activity during planning predicts the upcoming order of movements

To identify what content is represented in the underlying activity patterns during planning and production, we trained and tested an LDA classifier to distinguish between the neural patterns related to sequence identification, specifically of the planned and executed movement order, movement timing, and their unique combination^16,17^. All decoders were cross validated across six imaging runs and, for order and timing, across conditions, then converted into z scores (see Methods and materials).

We first tested whether ROI showed above chance decoding accuracy (Figure 3a, b). To do so, we obtained accuracy values for each of our three classifiers during preparation and production when considering all voxels within each ROI. We then assessed accuracy values in each region using one-tailed one-sample t-tests against a chance level of zero. All tests were Bonferroni corrected six times to account for classifier (3) and phase (2) in each pre-defined subcortical ROI. During preparation, we found above chance decoding of the upcoming sequence order in the left hippocampus (*t*(23) = 3.00, *p* = .018, *d* = 0.61). No evidence of significantly above-chance decoding accuracy could be found in the other ROIs (*p* > .264, *d* < 0.36). These results suggest that the left hippocampus is involved in the planning of the ordinal structure of movement sequences.

**Figure 3.**
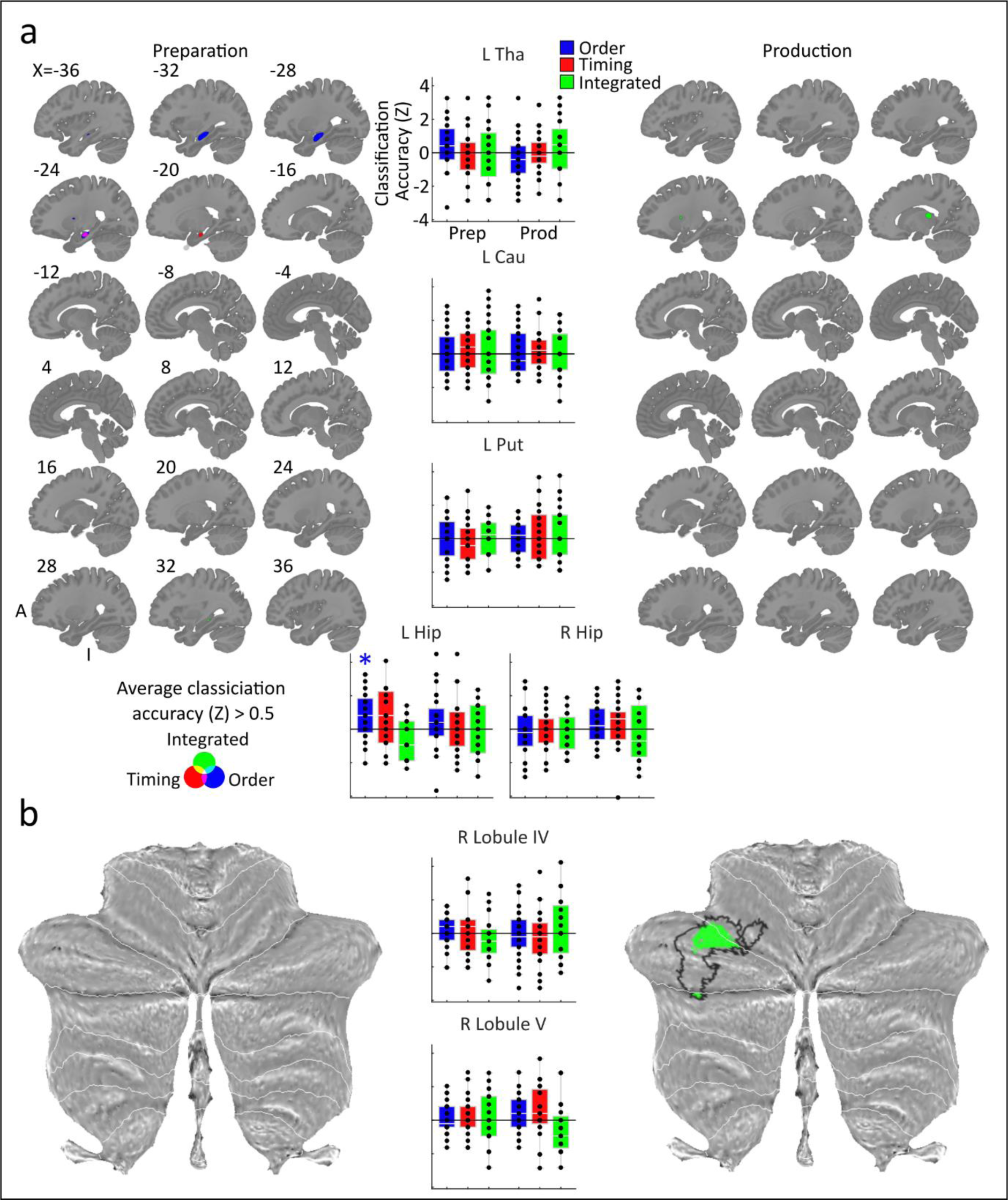
Linear discriminant analysis results. **a**, volumetric slices of each subcortical region displaying classification accuracy for each classifier during preparation and production, far left and right respectively. Centre shows classification accuracy when averaged across each subcortical region. * *p* < 0.05; one-sided t test against chance (Bonferroni-corrected for six comparisons within each ROI). Error bars represent standard error of the sample mean. **b**, as above, for cerebellum. Black outline indicates significant cluster. Tha, thalamus; Cau, caudate nucleus; Put, putamen; Hip, hippocampus.

Though considering all present voxels contained within a region is informative as to the general inclination of a region’s processing, it does not inform us as to which subregions may be driving decoding accuracy the most. This is especially true when it comes to subcortical regions such as the thalamus, which shows differing functional and anatomical connectivity throughout its various subregions and nuclei^30^. Therefore, we performed volumetric searchlight analyses using 160 voxel searchlights for each decoder, constrained to each subcortical ROI (Figure 3a, b). The accuracy value for each given searchlight was assigned to the centre voxel. We then identified significant clusters using a random effects analysis on the produced accuracy maps and applied SPM’s small-volume correction in a similar manner to the contrast activity cluster analysis. We Bonferroni corrected significance values for each cluster six times, to account for classifier (3) and phase (2). During preparation, we found a significant cluster for the order decoder in the anterior body of the left hippocampus (*t*(23) = 5.74, *pFWE* < .001, extent = 379, MNI coordinates: x=-32, y=-22, z=-11). No significant clusters were found during production, although prior to six-fold corrections a cluster for integrated sequence decoding was found in the superior thalamus during production (*t*(23) = 4.76, *pFWE* = .045, extent = 47, MNI coordinates: x=-18, y=-26, z=15). These findings reinforce that the hippocampus, particularly the anterior body, is involved in the planning of sequential movement order. Sequence timing and its non-linear integration with sequence order, however, do not seem to be represented in effector-related subcortical regions.

### Sequence-related activity patterns are not conserved across planning and execution

Given that our target subcortical regions show different activity levels alongside changes in informational content across preparation and production, we asked whether the sequence related patterns are maintained prior to and during movement. One possibility is that the patterns present during preparation undergo linear scaling to exceed a set threshold and initiate movement, where unintended initiation is thought to be prevented by suppression in corticospinal circuits^31^. Alternatively, these patterns may alter their geometries, meaning that they undergo a significant state change. The presence of scaled cortical neural patterns has been evidenced in single finger movements using fMRI^32^, whereas distinct neural patterns have been shown in multi-unit recordings of cortical populations^33^, however, subcortical pattern dynamics before and during sequence production are unknown. Accordingly, we assessed the within-sequence, across-phase Euclidean distance between conditions in our fMRI data across principal components using multi-dimensional scaling of the representational dissimilarity matrix. Since we saw substantial activity changes across all regions except for the caudate from preparation to production, the first principal component is set to represent the different levels of activity as it captures the dimension with the greatest variance across the conditions. Therefore, we quantified the Euclidean distance between each sequence’s position during preparation and production in a 2D space with the axes comprising of principal components two and three.

If one were to assume that activity patterns during production are simply upscaled versions of those during preparation, the hypothetical distance between patterns after removal of the first principal component should be zero. However, given a non-zero level of noise in our data, these distances are unlikely to equal zero which would bias our results away from indicating a maintained distribution. To control for this, we simulated fMRI data sets with the same conditions as our empirical data and generated preparation and production from either the same distribution (no switch) or different distributions (switch), whilst matching their signal to noise ratios to our regions of interest (Figure 4a). Additionally, we scaled this cross-phase distance by the average of distances within preparation and production (Figure 4b). We then performed one-tailed, one-sample *t*-tests against zero to identify significantly elevated Euclidean distance in each of our subcortical regions, Bonferroni corrected seven times for region (Figure 4c). We found significant elevation in the thalamus (*t*(23) = 6.06, *p* < .001, *d* = 1.24), caudate (*t*(23) = 3.20, *p* = .014, *d* = 0.65), putamen (*t*(23) = 3.12, *p* = .017, *d* = 0.64), hippocampus in the left (*t*(23) = 3.05, *p* = .020, *d* = 0.62) and right (*t*(23) = 3.93, *p* = .002, *d* = 0.80) hemispheres. No significant differences were found in CB lobule IV (*t*(23) = 0.37, *p* = 1.00, *d* = 0.08) and CB lobule V (*t*(23) = 2.05, *p* = .182, *d* = 0.42). These results suggest that all subcortical regions apart from CB show significantly higher cross-phase distances than a matched simulation which does not predict a change in distributions across phase, providing evidence for a substantial state change which alters sequence-specific neural patterns form prior to and during movement.

**Figure 4.**
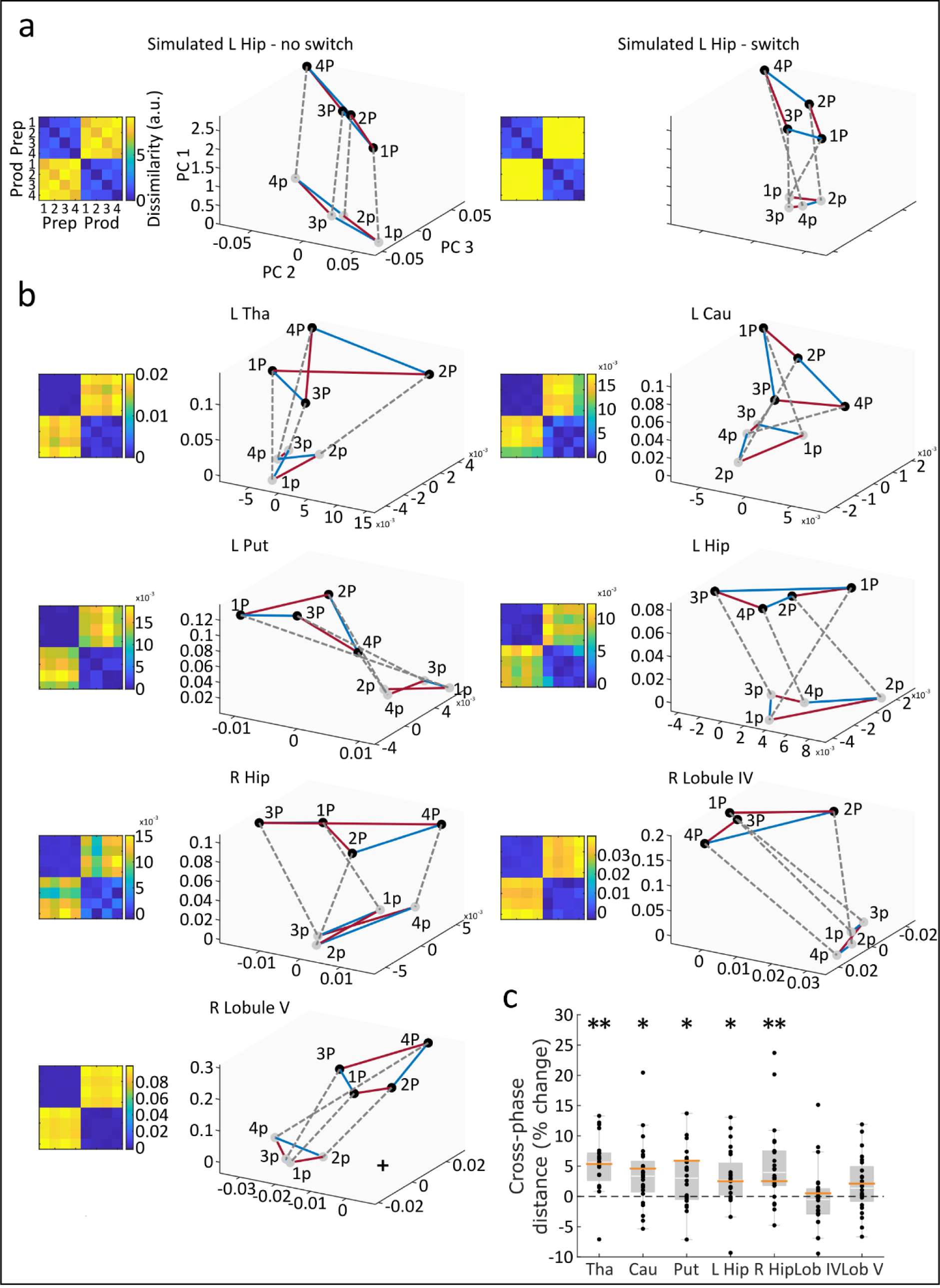
Simulated and empirical cross-phase Euclidean distances using multi-dimensional scaling. **a**, multi-dimensional scaling plots of simulated fMRI data along principal components 1, 2, and 3, showing four simulated sequence conditions across two phases. Red lines connect simulated sequences with different timings but the same order, whereas blue lines indicate sequences with different orders but the same timing. Dotted grey lines are drawn between the same sequence during preparation and production. Preparation and production were either generated using the same distribution (left panel; no switch) or different distributions (right panel; switch). **b**, multi-dimensional scaling plots of empirical data from target ROIs showing all four sequences during preparation and production. **c**, averaged Euclidean distance between preparation and production within sequences once PC1 is excluded, calculated relative to those same distances from the simulated no switch data with matched signal to noise ratios. This represents differences for all sequences between preparation and production which cannot be explained by differences in overall activity. Orange horizontal lines represent the same distances in the simulations when a switch is induced. * *p* < 0.05; ** *p* < 0.001; one-sided t test against simulated no distance with matched noise, 0 (Bonferroni-corrected for seven comparisons). Hip, hippocampus; Tha, thalamus; Cau, caudate nucleus; Put, putamen; Lob IV, lobule IV; Lob V, lobule V.

## Discussion

Acquiring a repertoire of skilled motor sequences has been linked to plasticity in subcortical motor-related brain areas – basal ganglia, the cerebellum, and, more recently, the hippocampus^9,27,34^. However, the contribution of these areas to higher-order action selection and lower-order movement control in skilled movements is debated^35–38^. Here, we examined the contribution of effector-related subcortical regions and the hippocampus to motor sequence control in the peri-movement phase (Figure 5). Further, we assessed whether the neural patterns during planning in those subcortical areas were simply a subthreshold versions of those during execution^31,39^, or qualitatively distinct neural states^33^. We found that both hippocampal and effector-related striatal regions showed increased activity during sequence planning. However, only the hippocampal activity was related to sequence-specific information, specifically predicting the sequence order shortly before each execution. In contrast, effector-related striatal, thalamic, and cerebellar regions increased activity during execution, yet lacked sequence-specific tuning, suggesting their role may be related to the control of lower-level kinematics of individual movement elements, rather than of a whole movement sequence. All regions, apart from the cerebellum, showed fundamental shifts in the sequence-related activity patterns from planning to execution, indicating that movement onset is associated with a fundamental state shift in subcortical areas rather than a scaling up of subthreshold patterns, similar to the previous results in the cortex^16,33^.

**Figure 5.**
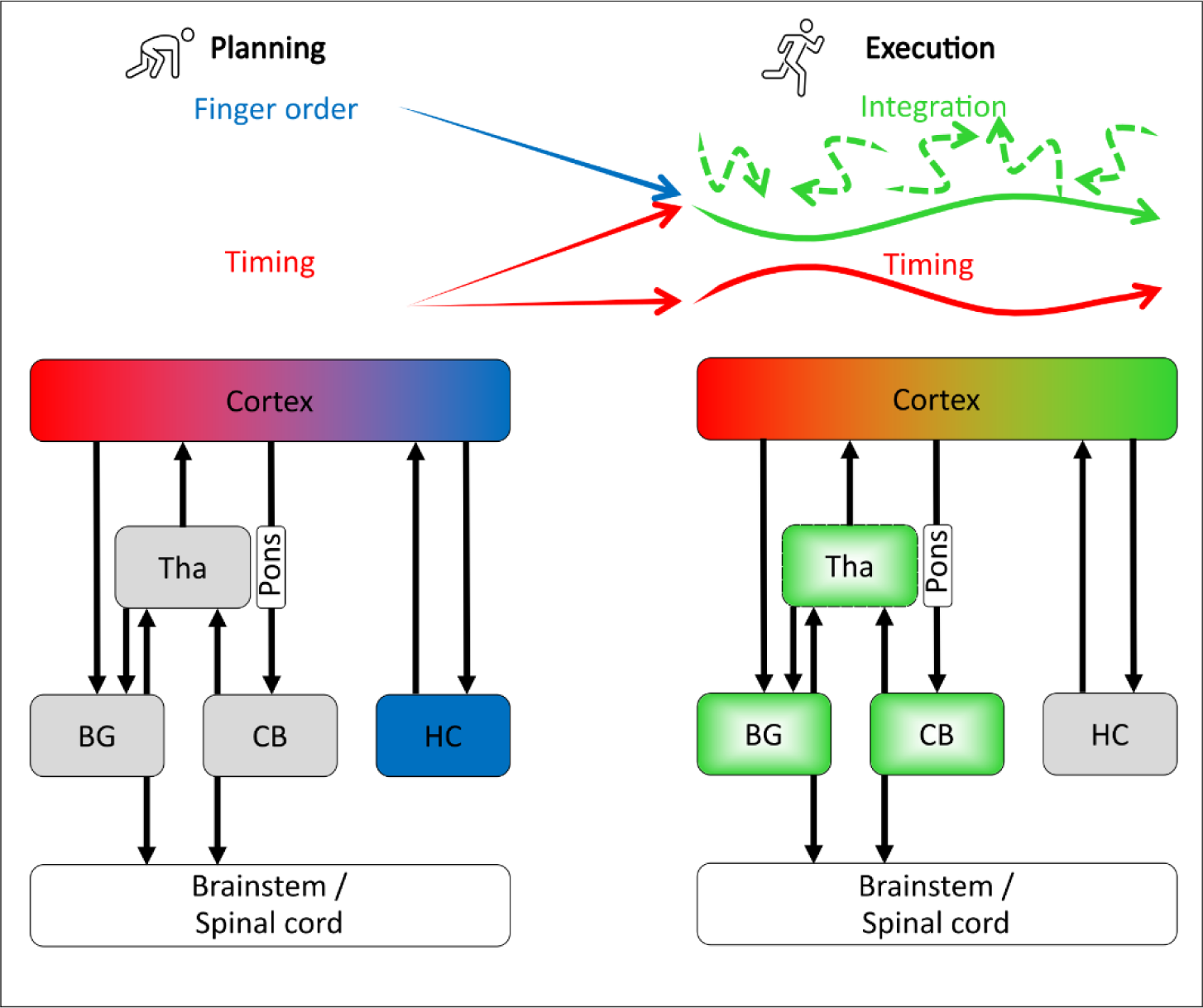
Schematic representation of sequence feature control during planning and execution. Higher level sequence features, order (blue) and timing (red), are defined by cortex and hippocampus during planning. During execution, the activity patterns in hippocampus, basal ganglia, and thalamus shift while the cortex integrates order and timing into low-level sequence specific trajectories (green), while maintaining higher-level independent movement timing. The green gradient fill in thalamus, basal ganglia, and cerebellum indicates regions that implement individual sequential elements. Tha, thalamus; BG, basal ganglia; CB, cerebellum; HC, hippocampus.

### The hippocampus pre-orders upcoming movements

The hippocampus has traditionally been associated with episodic recall in the non-motor domain and is key to the preservation of event order^40,41^. Here we show that the hippocampus is part of online motor control of a procedural skill which was trained over several days and produced without external guidance. Specifically, it is key to setting up the serial order of movements in an upcoming sequence from long-term memory (Figure 5). This is a high-order control function, as the same hippocampal patterns are retrieved regardless of the upcoming temporal structure (timing) of the sequence. These findings show that the hippocampal involvement goes beyond its contribution to the acquisition of motor skills, which has been reported in the context of sleep-related or wakeful consolidation during short rest periods interspersed with sequence learning^26,27^. Instead, it is participating in the rapid planning of a well-trained skill.

The hippocampus may retrieve the order of the upcoming motor sequence via competitive queueing of the upcoming movements^11,42,43^. This computational framework suggests that upcoming movements are pre-ordered in parallel prior to their serial execution during production in the parallel planning layer^44^ which has been confirmed experimentally^42,45^. Alternatively, the hippocampus may engage in serial order preplay^46^, which is temporally compressed approximately 20-fold relative to the acquired skill and is associated with skill consolidation^27^. How competitive queuing and preplay are related to each other is unclear. It is possible that they either measure the same underlying mechanism, e.g. pre-ordered sequence elements being related to incomplete replay sweeps, or constitute co-existing modes of serial order control across domains.

Our previous findings in the cortex suggest that the superior parietal lobule (SPL) also plays a role in the definition of movement order during planning^16^ and represents effector non-specific movement intentions ^47,48^. The SPL and hippocampus share extensive connections^49^ and interact closely during navigation tasks to convert world-centred into body-centred coordinates^50,51^. As such, interaction between the hippocampus and the SPL may serve to map the world-centred order retrieved by the hippocampus^42^ onto intrinsic reference frames^52^ prior to execution.

### Effector-relevant striatal, thalamic, and cerebellar activity lacks sequence-specific tuning

Despite robust activity increases, in particular during sequence execution, we found no evidence that effector-relevant contralateral striatal, thalamic, or ipsilateral cerebellar activity contained any sequence-specific information. These findings speak against both the hypothesized role of the striatum in higher-order action selection ^36,53–55^ or the integration of sequence features for kinematic control of the whole sequence as one motor program^56–58^. Human neuroimaging studies report movement concatenation-related activity in the striatum^59^, however our data suggests that these control signals are not sequence-specific^60^, but might instead be related to the process of concatenation itself. Further, multi-unit recordings in the dorsomedial striatum (caudate nucleus in humans) showed ramping activity which scales with temporal intervals between movements^61,62^, and is sensitive to the temporal structure of sequential movements^63^. However, we find no evidence that the striatum encodes differences in upcoming or ongoing temporal interval structure of sequences that are matched in structural complexity – neither during planning, nor during execution.

Recent results in rodent models showed that optogenetic lesions lead to kinematic deficits in overtrained sequence production^38^. Interestingly, this mapping breaks down when highly trained sequences share elements with other sequences the animal has been exposed to, e.g. lever presses appearing in a different order, with the skill becoming once-more cortically-dependent^64^. This is typical for many motor skills in humans, such as typing, handwriting and music production where elements are reused in different permutations, including highly overtrained sequences such as signing with one’s name or writing “Best wishes”. It is reflected in the factorial design of our study, which requires a well-trained recombination of movements into new sequences and shows no evidence of striatal tuning to these sequences. Thus, kinematic control in the striatum is likely associated with fused movements that are not broken up for flexible recombination – here, individual muscular commands associated with the different finger presses on the force transducer.

Cerebellar motor regions have traditionally been associated with low-level sensorimotor learning and online motor control, as evidenced by cerebellar ataxia^65,66^, adaptation to sensory perturbations^67^ and eye blink conditioning^35^. However, recent findings demonstrated the cerebellar involvement beyond short-timescales^68,69^. Transient perturbations of the deep cerebellar nuclei during planning resulted in mice making fewer correct directional licks and impacted subsequent movement direction whilst kinematics remained stable^37,68,69^. The interconnected ventrolateral thalamus, which constitutes an interface between striatum, cerebellum and cortex, has been shown to help maintain preparatory activity^70^ and control cortical dynamics from preparation to production for rapid and precise motor behaviour^71^. This extends the notion of the cerebellum as an online motor controller and the thalamus as a sensorimotor relay^72,73^. However, we did not observe any activity increase or sequence related patterns in ipsilateral effector-relevant cerebellar regions during motor sequence planning. This supports the classical findings that cerebellar motor regions contribute to the online control of individual movement elements during movement production, rather than the planning and execution of a whole sequence, e.g. planning “on the fly” once the production of a sequence has started^74,75^. Notably, sequence-related patterns could be found in cerebellar regions known to be interconnected with prefrontal areas (Crus I). This was only observed during motor execution, again emphasizing the role of the cerebellum in the online control of movements beyond effector-related regions.

Despite participants’ ability to fluently produce the target movement sequences from memory after two days of training, the movement kinematics for the upcoming sequence are not pre-planned. Rather, the order and timing of sequences is recombined trial-by-trial likely through communication with the cortex, which has shown a mechanism whereby the order and timing are planned and subsequently integrated during execution in the cortex^16^. This maintenance of separation up until the go cue likely affords the individual the ability to adjust movement plans should task demands suddenly change.

### Shift in sequence pattern activity across planning and execution

Motor planning in the cortex has been proposed to be a scaled down version of execution, held at bay by suppression in corticospinal circuits ^31,39^. More recently, however, planning activity in the motor cortex has been proposed to exist in an output-null dimension to execution, where increases and decreases in output from neurons projecting to the same source cause no net change in activity, but prepare the system to shift into the output-potent dimension ^33,76^, including for sequences of movements ^16^. Our current results suggest that this shift also takes place in multiple subcortical areas: striatum, thalamus, and hippocampus. Here, sequence-related patterns across planning and execution were significantly distinct across the peri-movement phases. This could not be explained by a simple upscaling through changes in activity and matched by our no-change simulations of the same dimensions that considered empirical noise levels in those areas. This could be driven by separate populations becoming active at different times around movement execution or, alternatively, changes of the patterns of the same neuronal pool.

### Implications for clinical disorders

Here we show that the hippocampus, but not the basal ganglia-cerebellar-thalamic loop is carrying sequence-related information during the planning of skilled movements produced from memory. Thus, breakdowns in movement sequence control in Parkinson’s Disease^77^ and Cerebellar Ataxia^78^ cannot be explained by deficits in the planning or execution of entire sequences. Instead, we hypothesize that they might be a result of the reduced capacity to concatenate movement elements or execute movement elements from the motor repertoire. In contrast, our results predict that Alzheimer’s Disease which is associated with a degeneration of the hippocampal formation and the medial temporal lobe, affects everyday movement sequence production such as handwriting and ideomotor action organization^79^ because of the inability to retrieve the correct order of the upcoming movement elements^43^. We hypothesize that this applies particularly to contexts where movement sequences need to be retrieved from memory flexibly in difference permutations. In humans, this applies to everyday skilled actions which utilise the same elements from the motor repertoire across different ordinal sequences, including overtrained sequences of movements, e.g. in typing, handwriting and tool use.

In summary, our results question traditional classifications of motor skill memory and control, along with their associated subcortical substrates. Research on skilled movement planning in diverse clinical populations is needed to inform appropriate rehabilitation interventions and support, e.g. by strategically boosting access to compensatory sequence planning mechanisms to enhance performance.

## Methods and materials

### Participants

24 participants (14 female and 10 male; mean age = 21.00 years, SD = 1.64 years) completed the three-day experiment, having met all behavioural and imaging requirements. All participants had no history of neurological disorders. 23 participants were right-handed with a mean Edinburgh Handedness Inventory (https://www.brainmapping.org/shared/Edinburgh.php; adapted from ^80^) score of 75.22 (SD = 20.97, range: 25-100), one was left-handed with a score of -70. Although our preregistration (www.osf.io/g64hv) stated that we would exclude left-handed individuals, this participant’s data did not differ qualitatively from the rest of the sample so were included. Data collected from an additional 17 participants were excluded for the following reasons: one due to unforeseen technical difficulties with the apparatus, one because of a corrupted functional scan, and 15 further did not reach target performance after two days of training. Target performance required an error rate < 20% (group mean = 6.54%, SD = 6.03%) and statistically distinct sequence timing structures that were maintained across different sequence orders; the full details regarding the exclusion criteria have been described elsewhere^16^. Participants were recruited either through a participation panel at Bangor University and awarded module credits, or through social media and given monetary reward at a standard rate. Individuals with professional musical qualifications were excluded from recruitment. All participants provided informed consent, including consent to data analysis and publication, through an online questionnaire hosted by Qualtrics. The Bangor University School of Psychology Ethics Committee approved this experiment and its procedures (ethics approval number 2019-16478).

### Apparatus

Two custom-built force transducer keyboards captured presses from all 10 fingers on the right and left hands of participants as they produced movement sequences. Each key had a small groove where participants placed their finger and could be adjusted for comfort according to the size of the participant’s hand. Below each key, a force transducer sampled force at 1000 Hz (Honeywell FS Series, with a range of up to 15N). The keys were not depressible. Force acquisition occurred in each trial from 500ms before sequence cue onset to the end of the production period in production trials, and the end of the false production period in No-Go trials. Traces from the right hand were baseline-corrected trial-by-trial using the first 500ms of acquisition (500ms before the sequence cue) and smoothed to a Gaussian window of 100ms. Button presses were recognised when a channel’s force exceeded a fixed threshold relative to baseline (2.5N for the first 8 participants and 1N for the subsequent 16 of 24 participants). The further apparatus used during the experiment have been described in a previous paper^16^.

### Behavioural task

Participants were trained to produce four five-finger sequences with defined timing structures, defined by inter-press intervals (IPIs), from memory in a delayed sequence production paradigm. Go trials began with an abstract fractal image (Sequence cue), which was associated with a sequence. Following the Sequence cue, a fixation cross was shown to allow participants to prepare the upcoming sequence. A black hand with a green background (Go cue) then appeared to cue sequence production from memory. Succeeding the Go cue, another fixation cross was presented. Feedback was then displayed to participants based on their performance during the preceding production period, finally followed by an inter-trial interval where a further fixation cross was displayed. During training, participants learned the sequences through repeated exposure to visually guided (Instructed) Go trials. These Instructed Go trials were functionally identical to From Memory Go trials, but featured a Go cue with a grey background and a red dot on the tip of each finger on the hand image, which moved from finger to finger in the target production order and in-pace with the target timing structure. No-Go trials had the same structure as Go trials, but the go cue did not appear following the preparatory fixation cross. Instead, the fixation cross continued to show for an extended period. This was succeeded by a further fixation cross, feedback, and ITI, as in Go trials.

The four five-finger target sequences consisted of permutations of two finger orders (Order 1 and 2) and two IPI orders (Timing 1 and 2) matched in finger occurrence and sequence duration. Sequence orders were generated randomly for each participant, making sure to avoid ascending and descending press triplets and any identical sequences. Furthermore, each participant’s trained sequences began with the same finger press to avoid differences in the first press driving the decoding of sequence identity during preparation ^81^. Timing structures were the same across participants, which consisted of four target IPI sequences as follows: 1200ms - 810ms - 350ms - 650ms (Timing 1), and 350ms - 1200ms - 650ms - 810ms (Timing 2).

The trial-by-trial feedback used a points-based scale, ranging from 0 to 10. Points were awarded based on initiation reaction time and temporal deviation from target timing. 0 points were awarded if the executed press order was incorrect. If the executed press order was correct, participants were awarded their earned timing points. In No-Go trials, 5 points were awarded if no press was made as instructed. Participants were presented with a feedback screen after each trial showing the number of points achieved in the current trial, as well as visual feedback on whether they pressed the correct finger at the correct time. A horizontal line was drawn across the centre of the screen, with four symbols displayed equidistantly along the line representing each of the five finger presses. Correct presses were indicated by an “X” symbol and incorrect presses were represented by a “–” symbol, for each respective sequence position. The vertical position of these symbols above (“too late”) or below (“too early”) the horizontal line was proportional to the participant’s timing of the respective press relative to target IPI. Using these cues, participants were able to adjust their performance online to ensure maximum accuracy of sequence production. During the first two days of training, auditory feedback in the form of successive rising tones corresponding to the number of points (0 - 10) was played alongside the visual feedback. Auditory feedback was absent during the fMRI session, to prevent any auditory processing driving decoding accuracy. All aspects of the behavioural task for this experiment are described in greater detail in a previous study^16^.

### Procedure

Training duration was consistent across participants and occurred across the first two days of the experiment over three distinct training stages. In the first training stage, 80% of all trials were instructed Go trials, and the remaining 20% were No-Go trials. During the second training stage, 40% of trials were instructed Go trials, 40% were from-memory Go trials, and 20% were No-Go trials. In the third and final stage of training, 80% of trials were from-memory Go trials, and 20% were No-Go trials. The third day took place inside of the MRI scanner and consisted of a short refresher stage, made up of the same proportion of trials as the second stage of training, during which T1 images were collected. Next, there was an fMRI stage consisting of six imaging runs, featuring 50% from-memory Go trials and 50% No-Go trials. In addition, before and after the last training stage, participants completed a synchronization task which has been described elsewhere^16^.

### MRI acquisition

Data were acquired using a 32-channel head coil in a Philips Ingenia Elition X 3T MRI scanner. T1 anatomical scans at a 0.937 x 0.937 x 1 resolution were acquired using MPRAGE, encoded in the anterior-posterior dimension with an FOV of 240 x 240 x 175 (A-P, R-L, F-H). For the functional data, T2*-weighted scans were collected across six runs of 230 volumes at a 2 mm isotropic resolution, with 60 slices at a thickness of 2 mm. The functional images were acquired at a TR of 2s, a TE of 35 ms, and a flip angle of 90°. These were obtained at a multiband factor of 2, in an interleaved odd-even EPI acquisition. To allow the stabilization of the magnetic field, four images were discarded at the beginning of each run. The whole brain of most participants was covered, except for the central prefrontal cortex, the anterior temporal lobe, and ventral parts of the cerebellum. Jitters were used within each trial during preparation periods, post-production fixation crosses, and ITIs, to give us a more accurate estimate of the Hemodynamic Response Function (HRF) by varying which part of the trial is sampled by each TR^82^.

### Pre-processing and first-level analysis

All fMRI pre-processing steps were completed using SPM12 (revision 7219) in MATLAB (The MathWorks). We applied slice timing correction using the first slice as a reference to interpolate all other slices to, ensuring analysis occurred on slices which represent the same time point. Realignment and unwarping were conducted using a weighted least-squares method correcting for head movements using a 6-parameter motion algorithm. A mean EPI was produced using SPM’s Imcalc function, wherein data acquired across all six runs were combined into a mean EPI image to be co-registered to the anatomical image. Mean EPIs were co-registered to anatomical images using SPM’s coreg function, and their alignment was checked and adjusted by hand to improve the alignment, if necessary. All EPI runs were then co-registered to the mean EPI image. For the GLM, regressors were defined for each sequence separately for both preparation and production, resulting in eight regressors of interest per run. Preparation- and production-related BOLD responses were independently modelled from No-Go and Go trials, respectively, to tease out activity from these brief trial phases despite the haemodynamic response lag^83^. The preparation regressor consisted of boxcar function starting at the onset of the Sequence cue in No-Go trials and lasting for the duration of the maximum possible preparation phase (2500ms). The production regressor consisted of a boxcar function starting at the onset of the first press with a fixed duration of 0 (constant impulse), to capture activity related to sequence initiation and extract sequence production-related activity from the first finger press that was matched across sequences within each participant. We aimed at capturing BOLD responses related to neuronal populations that become differentially active for different sequences^84^, for which a single estimate of sequence production has been used to successfully identify sequence representations in several previous fMRI studies^12,17,52,81,85^. Additionally, we included several regressors of no interest: (1) error trials (incorrect or premature presses during Go trials and presses during No-Go trials), which were modelled from sequence cue onset to the end of the ITI; (2) the preparation period in Go trials (1000- 2500 ms from Sequence cue); and (3) the temporal derivate of each regressor. The boxcar model was then convolved with the standard HRF. To remove the influence of movement-related artifacts, we used a weighted least-squares approach^86^. This resulted in us obtaining beta weight images for each of the eight conditions per scanning run, for all six runs. We further calculated the percent signal change during preparation and production relative to rest and compared each to a baseline of zero using two-tailed one-sample *t*-tests (Figure 2a). These tests were Bonferroni-corrected two times, to account for phase (2) within each pre-defined ROI. Additionally, we identified significant clusters of activity constrained to each ROI using a random effects analysis with an uncorrected threshold of *t*(23) > 3.48, *p* < 0.001 and a cluster-wise p value for the cluster of that size^87^.

### Subcortical and cerebellar regions of interest

We used Freesurfer’s automatic segmentation^88^ to segment subcortical regions of interest consisting of the thalamus, caudate nucleus, putamen, and hippocampus, from each participant’s T1 anatomical image. We then resliced each region’s mask into the same resolution as the functional images (2×2mm isotropic) and further masked them using functional activity. Only beta weights from voxels within these subcortical functional masks were extracted, constraining analyses to voxels belonging to subcortical regions of interest. Further, to assess the spatial organisation of multivariate patterns, we defined 160-voxel volumetric searchlights with a maximum radius of 6 mm in native space within each subcortical region of interest. LDA accuracy and RSA dissimilarity values (see Linear discriminant analysis and Representational similarity analysis sections) obtained were assigned to the centre voxel of the active searchlight.

For the CB, we used the SUIT cerebellar toolbox^89^ to segment the whole structure based on individual anatomical T1 images. We then resliced the lobular probabilistic cerebellar atlas^90^ into each participant’s native space to acquire masks for CB lobules IV and V. We then resliced these masks into functional resolution and further masked them using functional activity. Beta weights were extracted from these regions using these masks to constrain analyses to voxels of interest. Additionally, we defined a 160-voxel volumetric searchlight across the whole cerebellum using each participant’s individual anatomy. LDA accuracy and RSA dissimilarity values were assigned to the centre voxel of the active searchlight in a similar manner to the subcortical analysis. Classification accuracy and distance maps were subsequently resliced into SUIT space to display group results on a cerebellar surface flatmap^91^.

### Linear discriminant analysis

LDA was used to detect sequence-specific representations^16,17,42^, programmed in a custom-written MATLAB (The MathWorks) code. We extracted mean patterns and common voxel-by-voxel covariance matrices for each class from the training dataset (five of the six imaging runs), and then a gaussian linear discriminant classifier was used to distinguish between the same classes in the test dataset (the remaining imaging run). The factorized classification of finger order, timing, and integrated order and timing followed the previous approach^16,17^ and was performed on betas estimated from the sequence preparation and production periods independently. For the decoding of sequence order, the classifier was trained to distinguish between two sequences with different orders but matching timing across five runs and was then tested on two sequences with the same orders paired with a different timing in the remaining run. This classification was then cross-validated across runs and across training/test sequences, for a total of 12 cross-validation folds. For the decoding of sequence timing, the classifier was trained to distinguish between two sequences with differing timings paired with the same order and tested on two sequences with the same two timings paired with a different order and underwent the same cross-validation procedure. This method of training and testing the linear discriminant classifier allowed for identification of sequence feature representations that were transferrable across conditions they are paired with and therefore independent. The integrated classifier was trained to distinguish between sequences on five runs and then tested on the remaining run. Here, the mean activity for each timing (collapsed across two orders) and finger order (collapsed across two timings) condition within each run was subtracted from the overall activity for each run, separately^16,17^. This allowed for the measurement of residual activity patterns that were not explained by a linear combination of timing and order, rather spatiotemporal idiosyncrasies that would result in the generation of unique kinematics for the respective sequence that were not transferrable to others. We then trained our integrated classifier to distinguish between all four sequences based on these residual activity patterns across five imaging runs and then tested its accuracy on the remaining imaging run, cross-validated across runs.

We normalised classification accuracy by transforming to z scores, assuming a binomial distribution of the number of correct guesses^17^. We then tested these z scores against zero (chance level) in subcortical and cerebellar ROIs across participants for statistical analysis. Z-score transformed classification accuracy maps were subjected to a random effects analysis, similar to the RSA distance maps, with an uncorrected threshold of *t*(23) > 3.48, *p* < .001 and a cluster-wise p value for a cluster of that size^87^. Analyses that considered all voxels within respective regions of interest were also subjected to one-tailed one-sample *t* tests relative to zero, Bonferroni-corrected 6 times to account for phase (2) X classifier (3), to identify decoding accuracy above chance level. We also ran repeated measures ANOVAs on regional distance measures with factors of phase, classifier, and region to assess interaction effects and ran post-hoc pairwise comparisons to investigate significant interaction effects.

### Multi-dimensional scaling

We firstly pre-whitened beta weights prior to multi-dimensional scaling to increase the reliability of our distance measurements, using a regularised estimate of the overall noise-covariance matrix^92^. We then identified the first three principal components in our pre-whitened beta weights using classical multi-dimensional scaling of the variance-covariance matrix. This method plots all eight conditions in 3D space according to the first three eigenvectors (principal components) associated with the largest eigenvalues. Given that activity levels varied within regions across preparation and production (see Results), we measured the Euclidean distance between preparation and production within each sequence upon exclusion of the first principal component, since the largest eigenvalues are likely associated with differences in activity levels across phase. Hence, we calculated the Euclidean distance between points plotted in the 2D space represented by principal components one and two. However, given a non-zero level of noise in the data, the Euclidean distance between two points will be biased away from zero. We therefore simulated fMRI activity patterns with a matched number of voxels, sample size, and signal to noise ratio relative to the mean across participants of each ROI collected in the empirical data. We then calculated the cross-phase within-sequence Euclidean distances to provide baselines for the one-sample t-tests carried out in each region (Bonferroni corrected seven times for region). Additionally, these distances were scaled and normalised according to the average of the average distances between conditions during preparation and production respectively.

